# The synergistic antitumor effect of combined Anti-Human Epidermal Growth Factor Receptor 2 (HER2) antibody and Gamma Interferon therapy to Ab resistant breast cancer cells

**DOI:** 10.1101/536490

**Authors:** Toshihiko Gocho, Hiromichi Tsuchiya, Shotaro Kamijo, Yoshitaka Yamazaki, Yui Akita, Akiko Sasaki, Yuji Kiuchi

## Abstract

Anti-HER2 antibody is molecular targeted antibody for cancer therapy. Approximately 20% of breast cancers are characterized by overexpression of HER2 protein. However, the recurrence rate was 30% and the metastasis rate was 18% one year after treatment of Anti-HER2 antibody for HER2 positive breast cancer. The resistance to antibody treatment is a major problem for patients. We previously reported that Anti-HER2 antibody and Gamma Interferon (IFN-γ) combined therapy showed higher anti-tumor effect than usual therapy in vitro and in vivo mouse experiments.

In this study, we evaluated whether anti-HER2 antibody and IFN-γ combined therapy shows good synergistic effect against drug resistant HER2 positive breast cancer cells and higher antitumor effect than conventional clinical treatment. The resistant cell lines were made under the continuous presence of antibody until cell growth was not affected by the drug. We divided the resistant cells into the appropriate number of groups, which we and treated with anti-cancer therapy. We evaluated the antitumor effect for both in vitro study and in vivo mouse xenograft model prepared with the same immunogenicity. And we investigated the differences of immunofluorescence staining of CD8, Gr-1 and PDL-1 in tissues, especially related to immunity system.

The combined therapy showed significantly higher anti-tumor effect than other groups in vitro and in vivo experiments. The combined therapy affects anti-tumor immunity in this immunofluorescence experiment. Taken together, we showed the possibility that combined therapy could be an effective treatment option for anti-HER2 antibody resistant breast cancer, helping patients suffering from cancer progression after developing treatment resistance.

## Introduction

Human Epidermal growth factor 2 (HER2) belongs to the Her family and is a receptor with tyrosine kinase activity that affects cell proliferation, cell differentiation, apoptosis, and cell survival through signal transduction systems such as PI3K, MAPK, RAS, and SRC. The overexpression of HER2 is associated with carcinogenesis ^[1] [2] [3]^.

Overexpression and genetic amplification of HER2 is seen in 20 to 30% of breast cancer, 10 to 20% of stomach cancer and also other cancers (uterine cancer, head and neck cancer, and esophageal). The expression level of HER2 correlates with the malignancy of the cancer and is associated with poor prognosis ^[2] [4] [5]^.

Trastuzumab is a monoclonal antibody and is used for the treatment of breast cancer and gastric cancer. Trastuzumab is an anti-HER2 antibody which attaches to the ectodomain of HER2. The treatment outcome for HER2-positive cancer has improved since the appearance of Trastuzumab. ^[6][7]^.

Herceptin with chemotherapy is currently recommended as primary therapy for HER2-positive breast cancer and gastric cancer ^[8] [9]^. The Trastuzumab + Pertuzumab + chemotherapy is a standard therapy about HER2-positive unresectable breast cancer and distant metastasis breast cancer, and Herceptin + capecitabine or 5-fluorouracil plus cisplatin is the standard therapy for the HER2-positive gastric cancer, and that extends median overall survival: 13.8 months vs. 11.1 months; hazard ratio (HR) 0.74; 95 % confidence interval (CI), 0.60–0.91; *P* = 0.0046, vs chemotherapy alone ^[6]^

However, the cancer cells develop resistance tomedical treatment. In fact, nearly all progresses; HER2-positive metastatic patient eventually succumb to their disease. According to recent data, the 5-year survival rate of breast cancer is 5-10%, and the overall survival rate of Her2 positive gastric cancer treated with Herceptine + chemotherapy is only 13.8 months ^[6] [10]^. Although few studies on cancers that developresistance to antiHER2 antibody therapy have been conducted, the next best alternative therapy has not yet been established. ^[10] [11]^

Cancer immunotherapy has attracted attention in recent years. In fact, cancer immunotherapy was chosen as “breakthrough of the year” in 2013 in the journal *Science*. Check point inhibiters, including nivolumab and ipilimumab, has shown good results for unresectable cancers ^[12] [13] [14]^. New cancer immunological therapies are anticipated for the treatment of unresectable and metastatic cancer which are resistant to conventional therapy, with encouraging results from a variety of ongoing clinical trials^[15][16]^.

We showed that there was a very high antitumor effect for combination therapy of monoclonal antibody and IFN-γ using the experiment system that fixed the same immunogenicity for mouse, cancer cell line and the antibody. Briefly, we showed that the Ab and IFN-γ combined therapy ① acted in cancer cell and changes the malignancy itself, ② changed the signal transduction ③ affects the cell cycle and inhibits cell proliferation ④ accelerates CD8T cells cytotoxicity ⑤ decreases the number of Myeroid Dervied Suppressor Cells (MDSCs) which inhibits the antitumor effect and raises the immunoreactivity within the tumor tissue. ^[17]^

Therefore, we had the hypothesis that the anti-HER2 antibody and IFN-γ combined therapy might be useful for cancers that developed resistance to anti-HER2 antibody, so in this study, we evaluated whether Trastuzumab and IFN-γ combined therapy shows good synergy effect to drug resistant HER2 positive human breast cancer cells.

## Materials and Methods

### Cell lines and culture condition

#### H2N113

H2N113 tumor cell line was generated from female Balb/c MMTV-ErbB-2/neu transgenic mice spontaneous breast cancer. H2N113 cell lines were cultured in RPMI-1640 medium containing L-Glutamine (5ml/500ml) (Gibco Life Technologies, CA, USA), supplemented with 10% fetal bovine serum(Biosera, Kansas city, MO, USA), Sodium pyruvate(5ml/500ml) (Gibco Life Technologies, CA, USA), Glutamax (5ml/500ml) (Gibco Life Technologies, CA, USA) and MEM NEAA (5ml/500ml) (Gibco Life Technologies, CA, USA), Gibco penicillin-streptomycin liquid (5000 units/ml penicillin/5000 µg/ml streptomycin) (Gibco Life Technologies, CA, USA) at 37°C with 5% carbon dioxide in 95% air.

#### H2N113R

The resistant cell line (H2N113R) was made under the continuous presence of 7.16.4 antibody.

1 × 10^5^ H2N113 cells were seeded on 10 cm petri dish, and 7.16.4 antibody was added every 3 days and cells were passaged before getting confluent. The concentration of 7.16.4 antibody was gradually increased from 2 µg /ml to 20 µg /ml 40 µg/ml, 60 µg/ml, 80 µg/ml and 100 µg/ml finally.

#### Drugs

The 7.16.4 is used as the anti-mouse ErbB2 monoclonal antibody (Ab). The 7.16.4 Ab was purified from clone 7.16.4 hybridoma that was kindly provided by Mark Greene lab.

Mouse IFN-γ was purchased from PROSPEC (PROSPEC Protein Specialists, Rohovot, Israel). Docetaxel (DTX) was purchased from LC Laboratories (LC Laboratories, MA, USA). DTX was diluted with Dimethyl sulfoxide (DMSO). Anti-mouse PD-L1 was purchased from BioXcell. (mouse PD-L1 (B7-H1), BE0101, inVivoMab, New Hampshire, USA). About Ab and IFN-γ combined treatment, IFN-γ was added 30 minutes after Ab treatment in all experiments.

### Cell Growth in vitro proliferation assay

We set up three groups; control group, 7.16.4 group and 7.16.4+ IFN-γ group. The 1×10^5^ H2N113R cells were seeded to three 6-well plates containing normal culture media, which were divided equally into the three groups described above.. Eight hours after seeding the cells, drug treatment was initiated. The cell culture was analyzed at two separate times (3 and 7 days), the cell number was recorded, and the cell growth curve was developed. Nothing was added to the control group, while the 7.16.4 group was treated with 10 µg/ml of 7.16.4 Ab, 7.16.4+ IFN-γ group was treated with 10 µg/ml of 7.16.4 and 100 IU/ml of IFN-γ. Cell number was counted by automated cell counter (Bio-Rad Laboratories, Inc. Hercules, CA.).

### Mouse in vivo Experiment

Eight week-old female Balb/c mice were bought and bred for one week to adjust the environment for this experiment. 1×10^6^ H2N113R cells were injected subcutaneously into both sides of the back of the mice. We distributed the mice into 4 groups to adjust the tumor size between the four groups and started drug treatment at 14 days after tumor inoculation.

We treated PBS 100 µl to control group (n=9), 7.16.4 100 µg /100 µl PBS to 7.16.4 group (n=11), IFN-γ 10,000 IU/100 µl PBS to IFN-γ group (n=8). 7.16.4 Ab 100 µg /100 µl PBS and IFN-γ 10,000 IU/100 µl PBS to 7.16.4+ IFN-γ combined therapy group (n=11). Drugs were delivered by intraperitoneal injection. In the combined therapy group, IFN-γ was added 30 minutes after 7.16.4 injection. The drugs were given three times a week. Tumor size was measured three times a week with a digital caliper carbon fiber (19978, Sink, Niigata, Japan) and calculated using a simple algorithm (length × width × height).

### Mouse in vivo comparative experiment with chemotherapy

We did the same in vivo experiment using H2N113R cells. We divided the cells into 6 groups.

We treated PBS 100 µl to control group (n=5), 7.16.4 100 µg /100 µl PBS to 7.16.4 group (n=8), DTX 100 µg /100 µl DMSO to DTX group (n=7), 7.16.4 100 µg /100 µl PBS and IFN-γ 10,000 IU/100 µl PBS and DTX 100 µg /100 µl DMSO to 7.16.4+ IFN-γ + DTX combined therapy group (n=10), anti-PD-L1 antibody 100 µg /100 µl PBS to aPD-L1 group (n=7), 7.16.4 100 µg /100 µl PBS and IFN-γ 10,000 IU/100 µl PBS and aPD-L1 antibody 100 µg /100 µl PBS to 7.16.4+ IFN-γ+aPD-L1 combined therapy group (n=9). Drugs were delivered by intraperitoneal injection. In the combined therapy group, IFN-γ was added 30 minutes after 7.16.4 injection. Docetaxel and aPD-L1 were given three times a week. The others were performed three times a week. Tumor size was measured three times a week with a digital caliper carbon fiber (19978, Sink, Niigata, Japan) and calculated using a simple algorithm (length × width × height). Tumors were removed from the mice on day 21 of the drug treatment. Specimens were fixed in 10% formaldehyde for 24 hours.

### Fluorescent immunohistochemistry

Tumors were removed from the mice on day 16 of the drug treatment from mice in vivo experiment and from the mice on day 21 of the drug treatment from mice in vivo experiment with chemotherapy. Specimens were fixed in 10% formaldehyde for 24 hours.

Tumors specimens were cut at a thickness of 3 µm and fixed in 10% formaldehyde. Sections were stained by H. & E. Immunostaining was performed by incubating these sections again in 0.3% H_2_O_2_ for 5 minutes to remove endogenous peroxidase activity and then incubated in the Dako nonspecific blocking reagent for 5 minutes. These specimens were incubated with a CD8 (1:100) (LEAF(tm)Purified anti-mouse CD8a, 100715, BioLegend^®^,CA, USA), Gr-1 (1:100) (Purified anti-mouse Ly-6G/Ly-6c, 108401, BioLegend^®^,CA, USA), PDL-1 (1:1000) (mouse PDL-1 (B7-H1), BE0101, inVivoMab, New Hampshire, USA) for 1 hour at room temperature, followed by a secondary, TRITC-conjugated anti-rabbit IgG (A21428, Life Technologies, USA) and bisbenzimide H33342 (DojinDO Molecular Technologies, Inc., Kumamoto, Japan) in a humid chamber at 37°C for 30 min. The images were captured on microscope with BZ-X800 (Keyence, Osaka, Japan) and were quantitated with Hybrid Cell Count BZ-H4C software (Keyence, Osaka, Japan)

### Statistical analysis

Statistical analysis was carried out using YSAT 2013 (Igakutosho-shuppan Ltd., Toda, Japan). Differences in mean values were statistically analyzed using non-repeated measures ANOVA, followed by Dunnett’s test or the Student-Newman-Keuls test. Tumor volume differences between the Control and 7.16.4 + IFN-γ groups were statistically analyzed using unpaired Student’s t-test. Tumor volume differences between the Control and 7.16.4 + IFN-γ + aPD-L1 groups were statistically analyzed using unpaired Student’s t-test, with significance level set at p < 0.05.

## Results

### Combined 7.16.4 + IFN-γ therapy suppressed the H2N113R Cell Growth in vitro proliferation assay

The average cell number on day 3 was 4.40 × 10^5^ cells and that of day 7 was 67.6 × 10^5^ cells of control group.

The average cell number on day 3 was 3.80 × 10^5^ cells and that of day 7 was 61.8 × 10^5^ cells of 7.16.4 group.

The average cell number on day 3 was 3.04 × 10^5^ cells and that of day 7 was 49.5 × 10^5^ cells of 7.16.4 + IFN-γ group.

On day 7, the percentage of the average cell number of 7.16.4 + IFN-γ group was 73% of that of control group, whereas that of the7.16.4 group was 91% of the control group. 7.16.4 + IFN-γ combined treatment significantly decreased the cell number compared to control group (P < 0.05), whereas 7.16.4 group showed no significant difference to control group. (Figure1)

**Figure 1.**
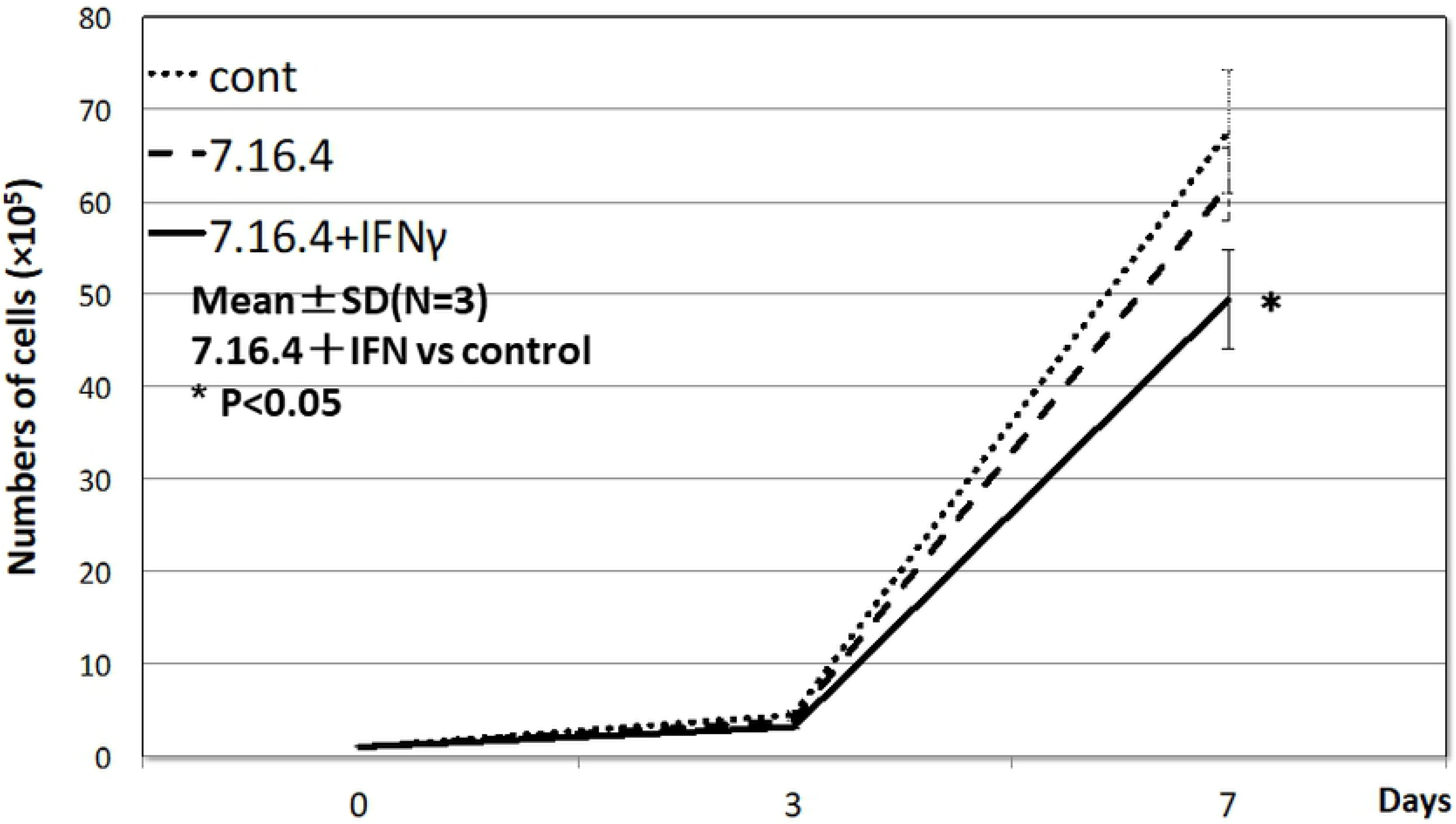
7.16.4 + IFN-γ therapy suppressed the H2N113R Cell Growth in vitro proliferation assay. The 1×10^5^ H2N113R cells were seeded to 6 well plate: Control,7.16.4 (10 µg/ml), 7.16.4 + IFN-γ (10 µg/ml + 100 IU/ml). Treatment was started after 8 hours from seeded and cultured 7 days. The cell numbers were counted on day 3 and 7 and the cell growth curve was made. Cell number was counted by automated cell counter (Bio-Rad Laboratories, Inc. Hercules, CA). 7.16.4 + IFN-γ combined treatment significantly decreased the cell number compared to control group (P < 0.05), whereas 7.16.4 group showed no significant difference to control group. The curves show the mean + standard deviation of the number of cells from independent 3 culture experiments. (Student’s t-test *p < 0.05 vs. control) (n=3)

H2N113R is not suppressed by 7.16.4 antibody. It means that we showed that H2N113R acquired the resistant ability for 7.16.4 antibody treatment after continuous presence of antibody, and we can use this cell line for antibody resistant cell line.

And 7.16.4 + IFN-γ combined treatment significantly decreased the cell number. This means that somehow the combined therapy effectively suppresses tumor growth in vitro without the effect of immune cells.

### 7.16.4 + IFN-γ therapy showed the antitumor effect against resistant H2N113R cell in mice Xenograft model

We set in vivo mice Xenograft experiment to investigate antitumor effects of 7.16.4 + IFN-γ under the condition of the effective immunity including the effect of antibody dependent cellular cytotoxicity (ADCC) and Complement-Dependent Cytotoxicity (CDC). We used the same immunogenicity for all stuff to investigate the real immunoreaction without immune response to invasion by foreign substances. For example, we used the Balb/c mice, H2N113R cells that is from Balb/c mice, 7.16.4 that is mouse IgG, mouse IFN-γ and mouse anti-PD-L1 antibody.

7.16.4 + IFN-γ treatment significantly suppressed tumor volumes compared with the other groups from day 7 until the end (Figure2). 7.16.4 + IFN-γ treatment showed continuous tumor regression by day 16, whereas treatment with 7.16.4 alone showed the tendency of tumor regression but not significantly tumor volume suppression compared with the other groups. 7.16.4 + IFN-γ treatment significantly decreased tumor volumes compared with Ab alone after day 7. On the other hand, IFN-γ alone didn’t show any anti-tumor effect, but rather, the tumor volume actually increased.

**Figure 2.**
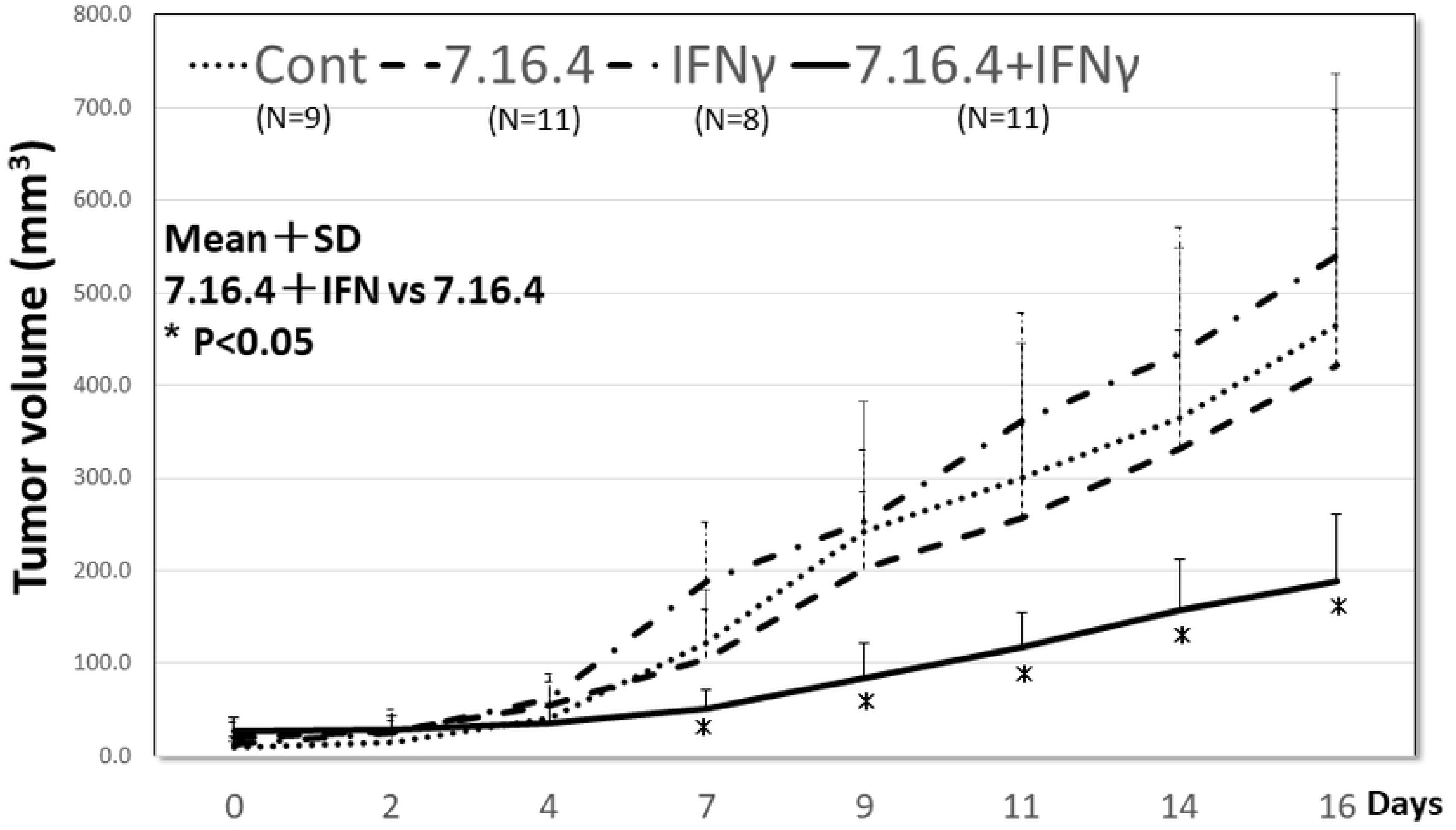
7.16.4 + IFN-γ therapy showed the antitumor effect against resistant H2N113R cell in mice Xenograft model. Balb/c mice were used. 1×10^6^ H2N113R cells were injected subcutaneously into both sides of the back of each mouse. We started drug treatment at day 14 post-injection (day 0 in the figure) after dividing into 4 groups: control (PBS 100 µl), 7.16.4(100 µg/100 µl PBS), IFN-γ (10,000 IU/100 µl PBS), 7.16.4 + IFN-γ (100 µg/100 µl PBS + 10,000 IU/100 µl). Tumor volumes were calculated as length × width × height. Measurements and drug treatments were performed three times a week. 7.16.4 + IFN-γ treatment significantly suppressed tumor volumes compared with the other groups from day7 until the end. (Student’s t-test *p < 0.05 vs. 7.16.4) The curve shows the mean + standard deviation of the volume of the tumors; the numbers of tumors shown as below (Cont: n=9, 7.16.4: n=11, IFN-γ: n=8, 7.16.4 + IFN-γ: n=10). (Dannet test, *p < 0.05 vs. control)

### 7.16.4 + IFN-γ therapy showed the promising potential to be applied to antitumor treatment against antibody treatment resistant tumors in clinical

We did the similar mice Xenograft experiment using 6 groups this time.

We selected the drugs the drugs refer to the clinical use.

7.16.4 + IFN-γ + aPD-L1 treatment showed the most tumor suppression compared with the other groups on day7 and from day12 to the end. (Figure3) (Bonferroni test, *p < 0.05, **p < 0.01 vs. control). Treatment with 7.16.4 + IFN-γ +DTX or aPD-L1 significantly suppressed tumor volumes compared with the control, aPD-L1, DTX and 7.16.4 alone treatment (*p < 0.05). On the other hand, treatment with aPD-L1, DTX and 7.16.4 alone didn’t show any significant difference in tumor suppression. This means that 7.16.4 + IFN-γ therapy is a key factor to treat antibody resistant tumor.

**Figure 3.**
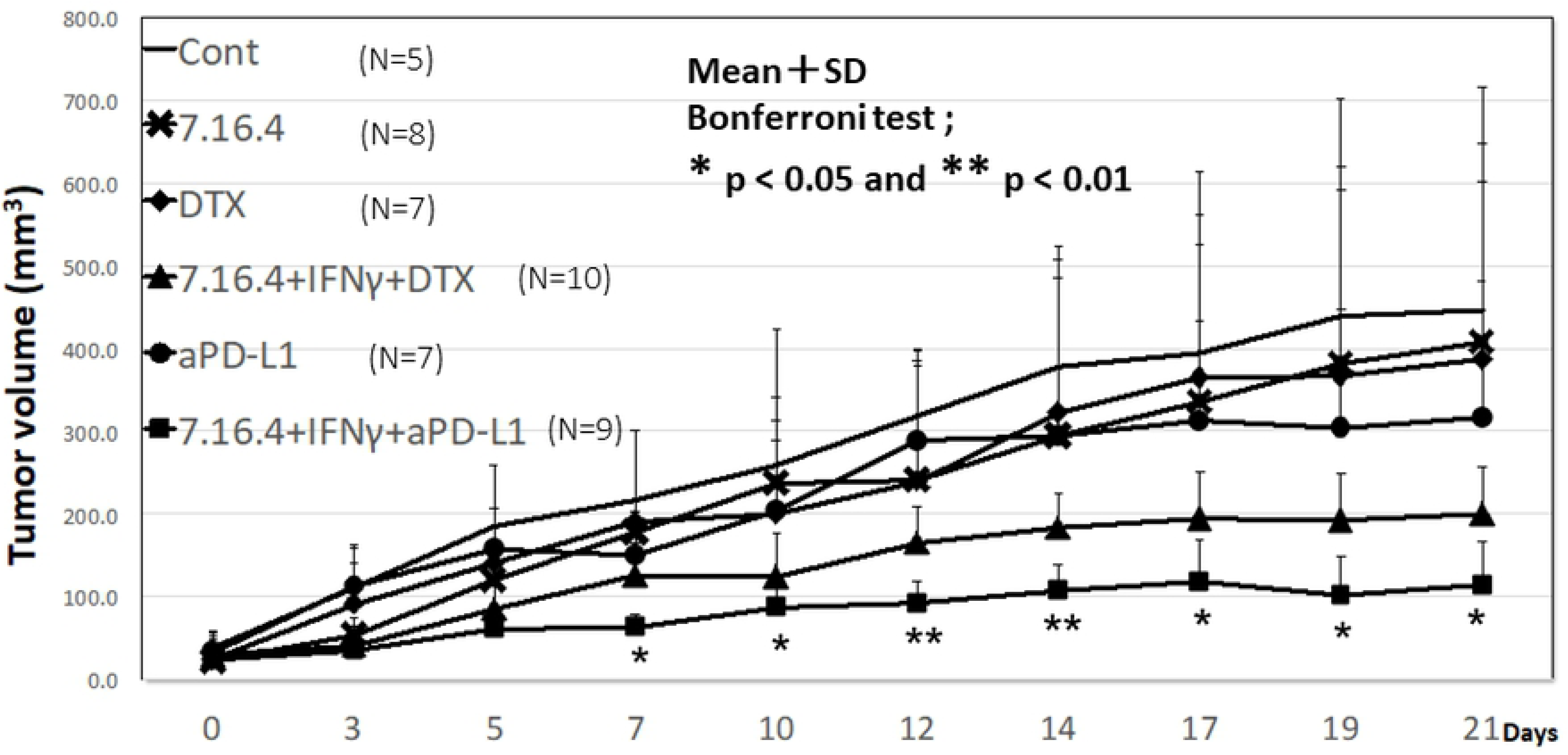
7.16.4 + IFN-γ therapy showed the promising potential to be applied to antitumor treatment against antibody treatment resistant tumors in clinical use. Balb/c mice were used. 1×10^6^ H2N113R cells were injected subcutaneously into both sides of the back of each mouse. We started drug treatment at day 14 post-injection (day 0 in the figure) after dividing into 6 groups: control (PBS 100 µl), 7.16.4 (100 µg/100 µl PBS), DTX (DTX 100 µg/100 µl DMSO), 7.16.4 + IFN-γ + DTX (100 µg/100 µl 7.16.4/PBS + 10,000 IU/100 µl IFN-γ/PBS + 100 µg/100 µl DTX/DMSO), aPD-L1 (aPD-L1 100 µg/100 µl PBS), 7.16.4 + IFN-γ + aPD-L1 (100 µg/100 µl 7.16.4/PBS + 10,000 IU/100 µl IFN-γ/PBS + 100 µg/100 µl aPD-L1/PBS). Tumor volumes were calculated as length × width × height. Measurements and drug treatments were performed three times a week. The curve shows the mean + standard deviation of the volume of the tumors; the numbers of tumors shown as below (Cont: n=5, 7.16.4: n=8, DTX: n=7, 7.16.4 + IFN-γ + DTX: n=10, aPD-L1: n=7,7.16.4 + IFN-γ + aPD-L1: n=9). 7.16.4 + IFN-γ + aPD-L1 treatment showed the most tumor suppression compared with the other groups on day 7 and from day 12 to the end significantly. (Bonferroni test, *p < 0.05, **p < 0.01 vs. control) Treatment with 7.16.4 + IFN-γ +DTX or aPD-L1 significantly suppressed tumor volumes compared with the control, aPD-L1, DTX and 7.16.4 alone treatment. (Bonferroni test, *p < 0.05, **p < 0.01 vs control)

### Histopathological examination from mice Xenograft model

Hematoxylin / Eosin (H&E) staining was performed to observe the differences in histopathological changes in tumor tissue, depending on the different treatment groups. The images were captured on microscope with BZ-X800 (Keyence, Osaka, Japan) and were quantitated with Hybrid Cell Count BZ-H4C software (Keyence, Osaka, Japan).

In the 7.16.4 + IFN-γ group, the tumor was characterized by atrophy? vacuolization 、 and interstitial lymphocyte proliferation. However, in the other groups, a large number of tumors had a swollen appearance. (Figure4)

**Figure 4.**
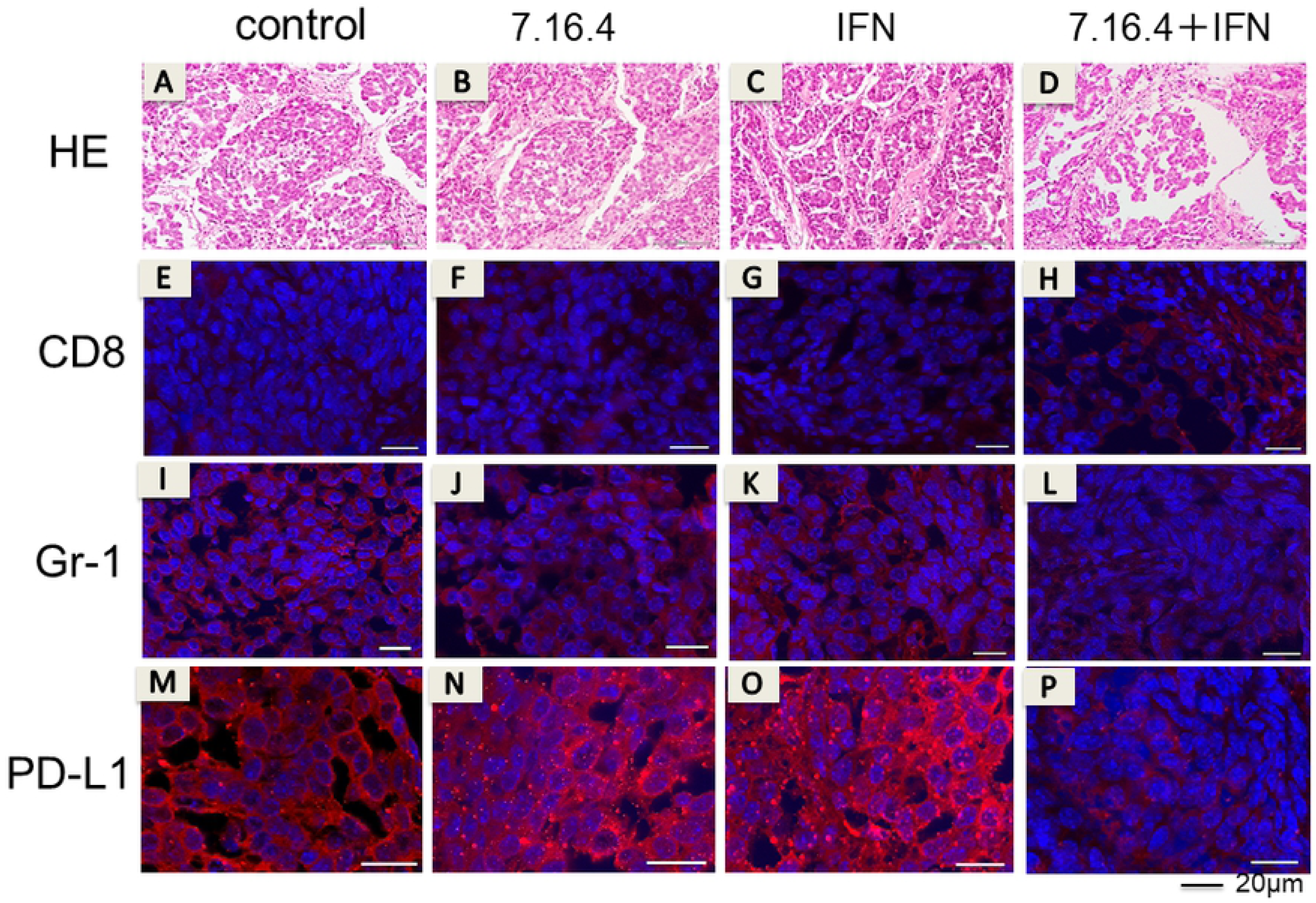
Histopathological examination from mice Xenograft model. Hematoxylin / Eosin (H&E) staining was performed to observe the differences of histopathological changes in tumor tissue depends on the different treatment groups. Tumors were removed from the mice on day 16 of the drug treatment from mice in vivo experiment. Images were captured on microscope with BZ-X800 (Keyence, Osaka, Japan) and were quantitated with Hybrid Cell Count BZ-H4C software (Keyence, Osaka, Japan). In the 7.16.4 + IFN-γ group, the tumor was atrophy, vacuolisetion, and interstitial lymphocyte proliferation. In the other group, a large number of tumors swoolen. The fluorescence intensity of the protein expression levels of CD8 of 7.16.4 + IFN-γ groups were significantly greater that of the other groups (p<0.05). And 7.16.4 + IFN-γ groups significantly reduced the intensity of Gr-1 and PD-L1 expression than the other groups (p<0.05). There was no significant difference in the expression level of CD8 and Gr-1 between control, 7.16.4 and IFN-γ groups. Compared with control group, PD-L1 expression significantly greater in mAb and IFN-γ alone groups (p<0.05). (Student T-test)

To investigate the immunity differences through each treatment for resistant breast cancer, the expression of the surface antigen proteins, CD8, Gr-1, PD-L1 was stained by immunofluorescence staining.

The fluorescence intensity of the protein expression levels of CD8 of 7.16.4 + IFN-γ groups were significantly greater that of the other groups (p<0.05). And 7.16.4 + IFN-γ groups significantly reduced the intensity of Gr-1 and PD-L1 expression than the other groups (p<0.05). There was no significant difference in the expression level of CD8 and Gr-1 between control, 7.16.4 and IFN-γ groups.

Compared with control group, PD-L1 expression significantly greater in mAb and IFN-γ alone groups (p<0.05) (Student T-test).

### Histopathological examination in clinical treatment

In the 7.16.4 + IFN-γ + DTX and 7.16.4 + IFN-γ + aPDL1 groups, the tumor was characterized by atrophy and interstitial lymphocyte proliferation. (Figure5)

**Figure 5.**
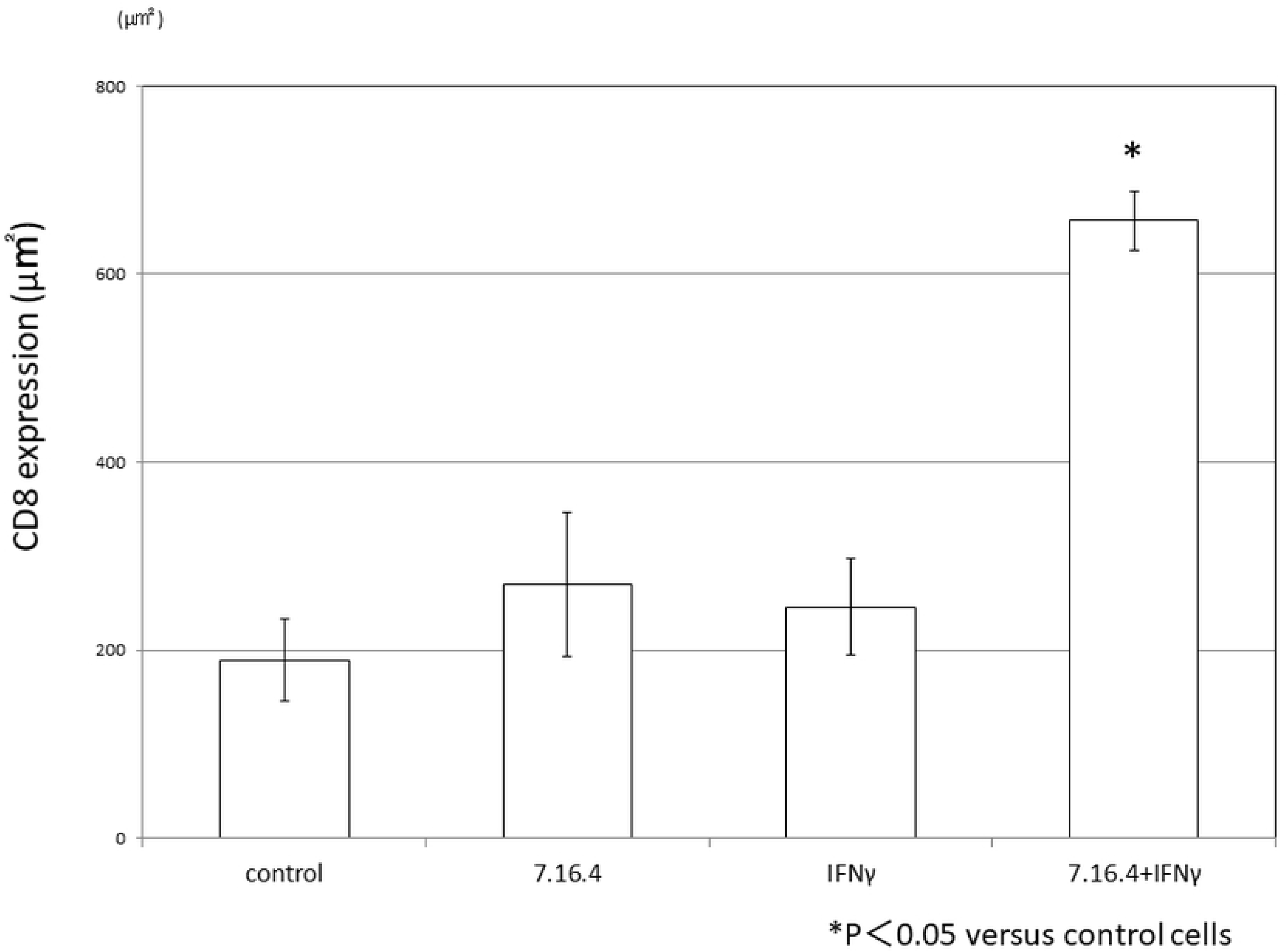
Histopathological examination in clinical treatment. H & E stain and the fluoresence intensity of CD8, Gr-1, and PD-L1 was performed and observed same as figure4. In the 7.16.4 + IFN-γ + DTX and 7.16.4 + IFN-γ + aPDL1 groups, the tumor was atrophy and interstitial lymphocyte proliferation. CD8 and expression was significantly higher in 7.16.4 + IFN-γ + aPD-L1 group (p<0.05) than the other groups. The Gr-1 expression was significantly lower in 7.16.4 + IFN-γ + aPD-L1 and 7.16.4 + IFN-γ + DTX group (p<0.05). In DTX alone group the PD-L1 expression significantly lower. And the PD-L1 expression in 7.16.4 + IFN-γ + PD-L1 group was significantly increased than control group. (p<0.05). (Student T-test)

**Fig. 6.**
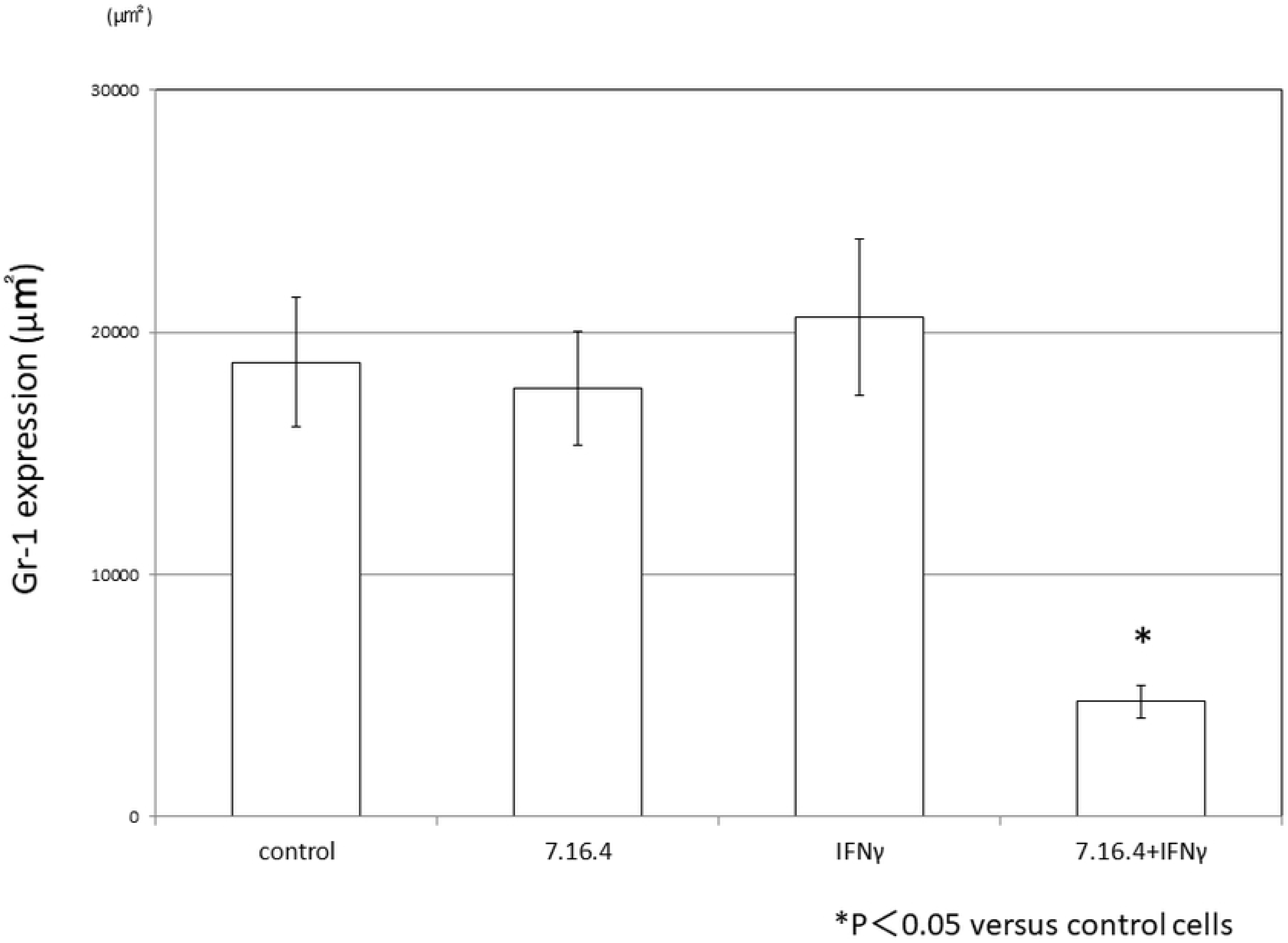

**Fig. 7.**
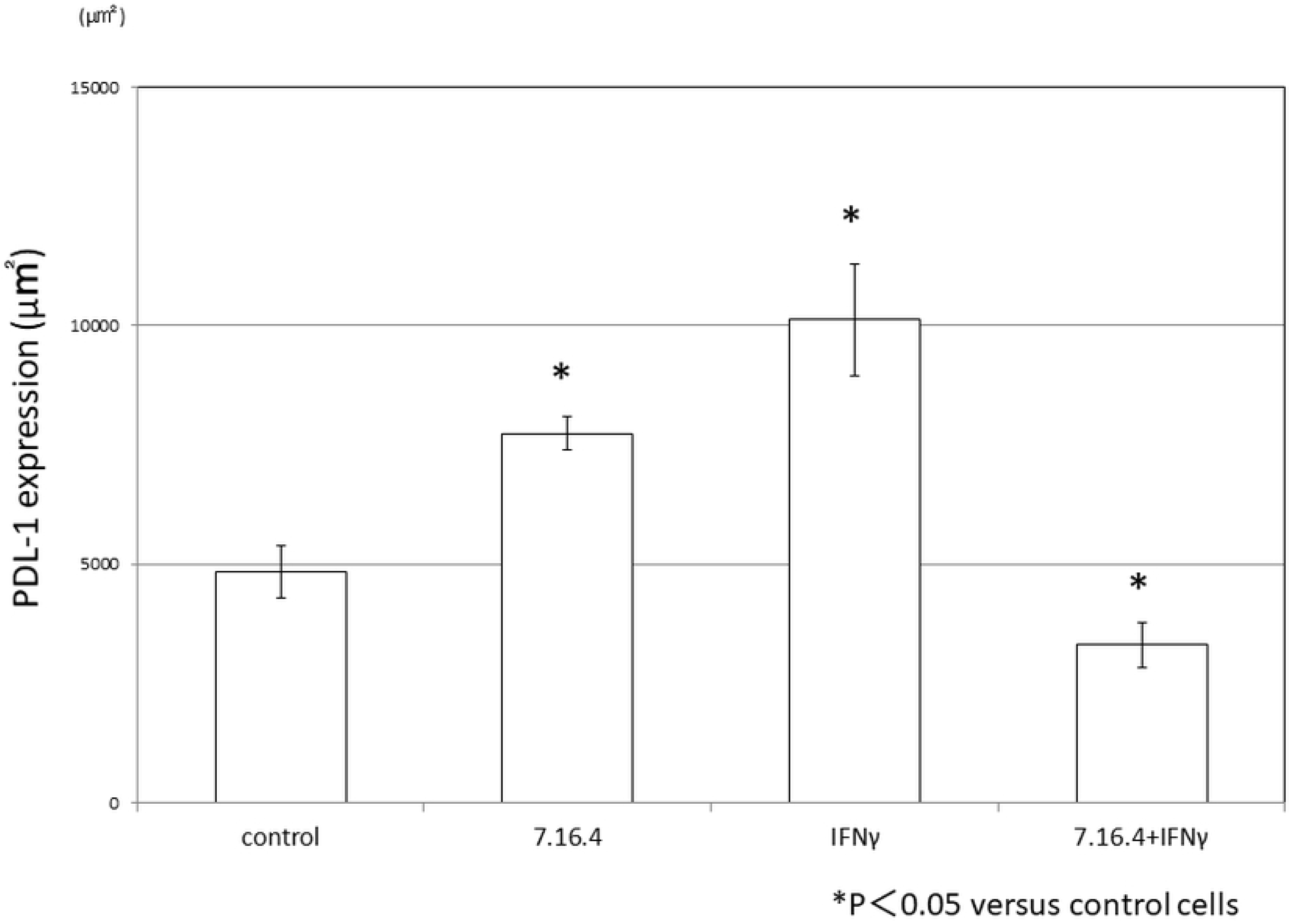

**Fig. 8.**
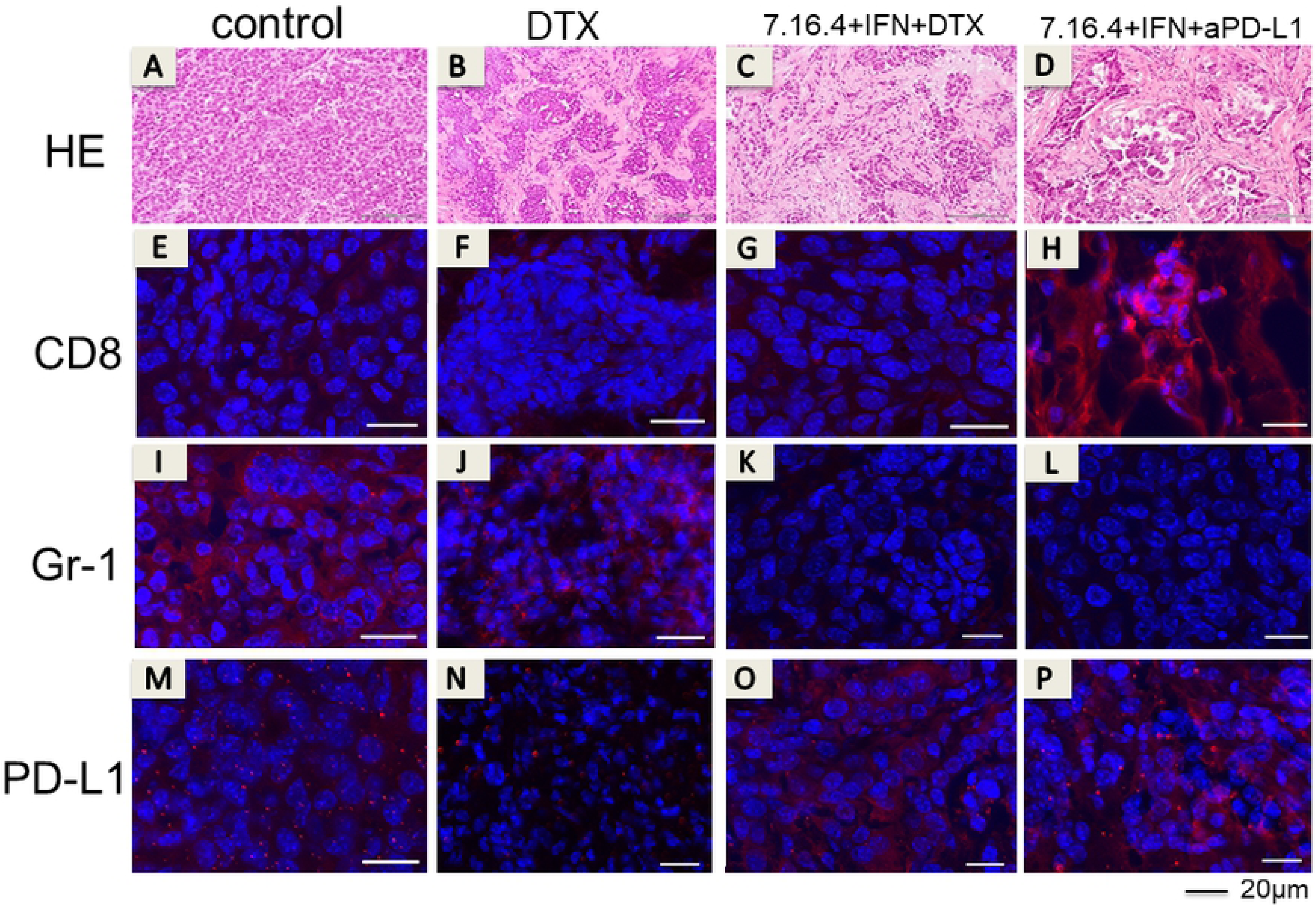

**Fig. 9.**
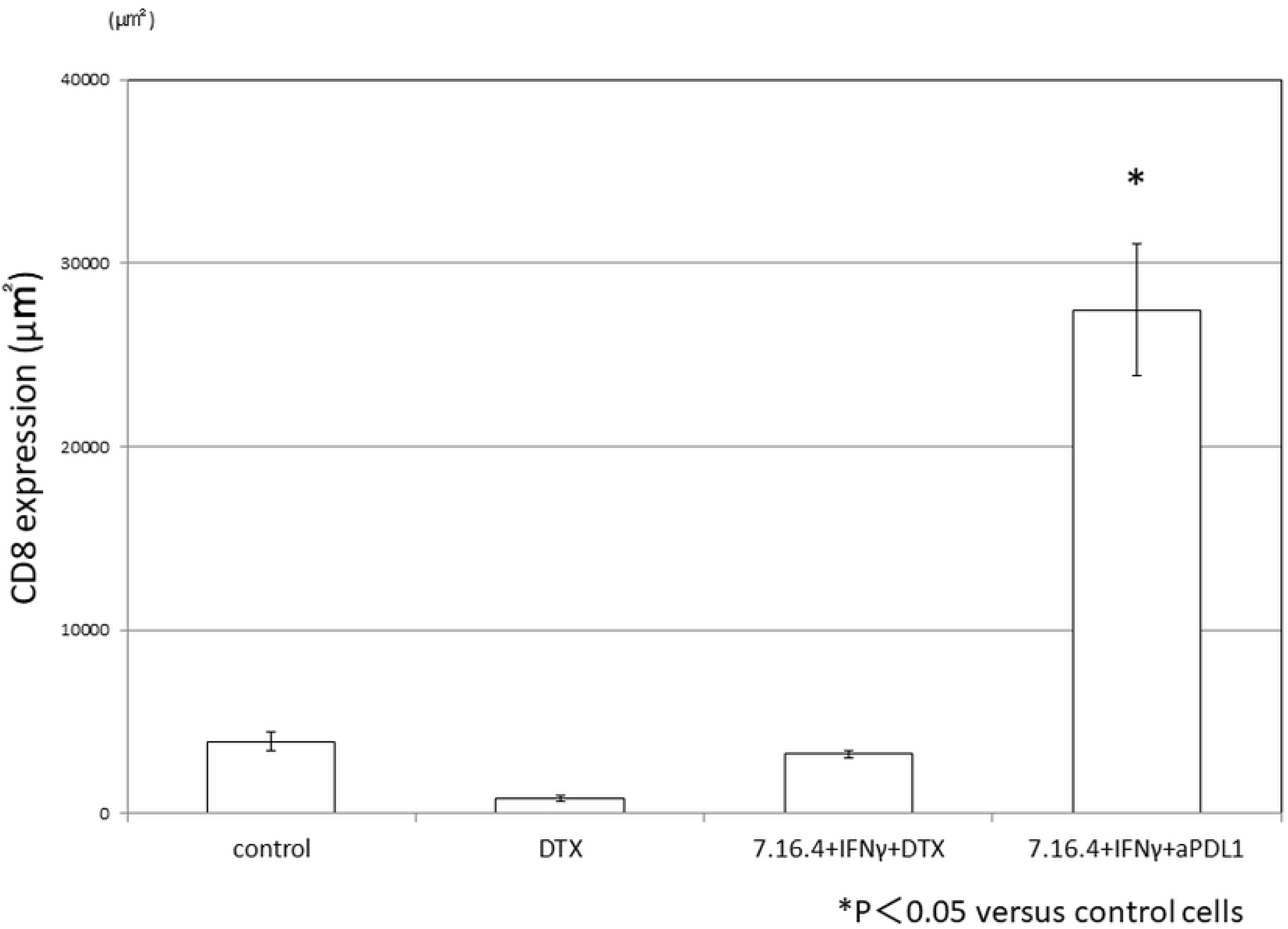

**Fig. 10.**
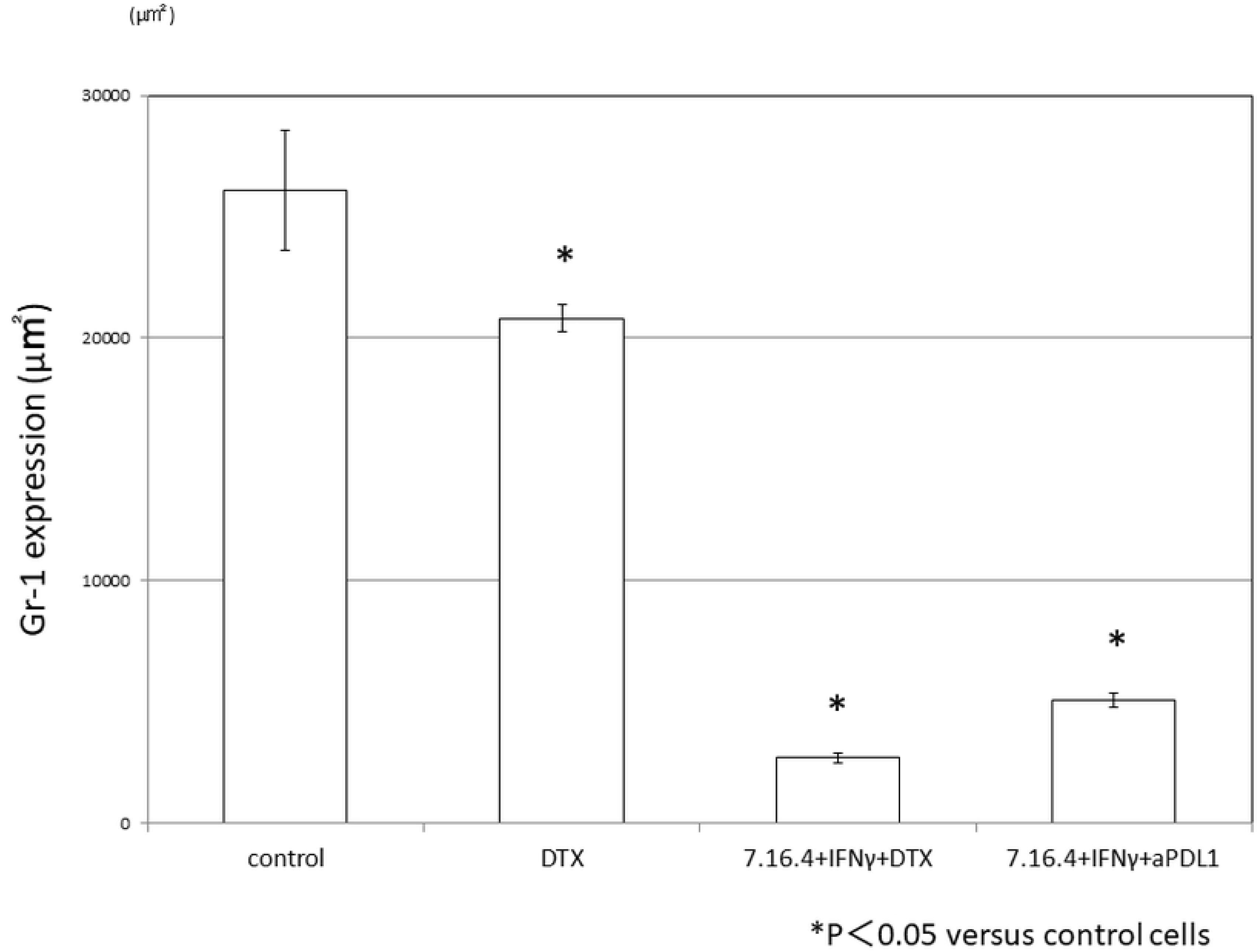

**Fig. 11.**
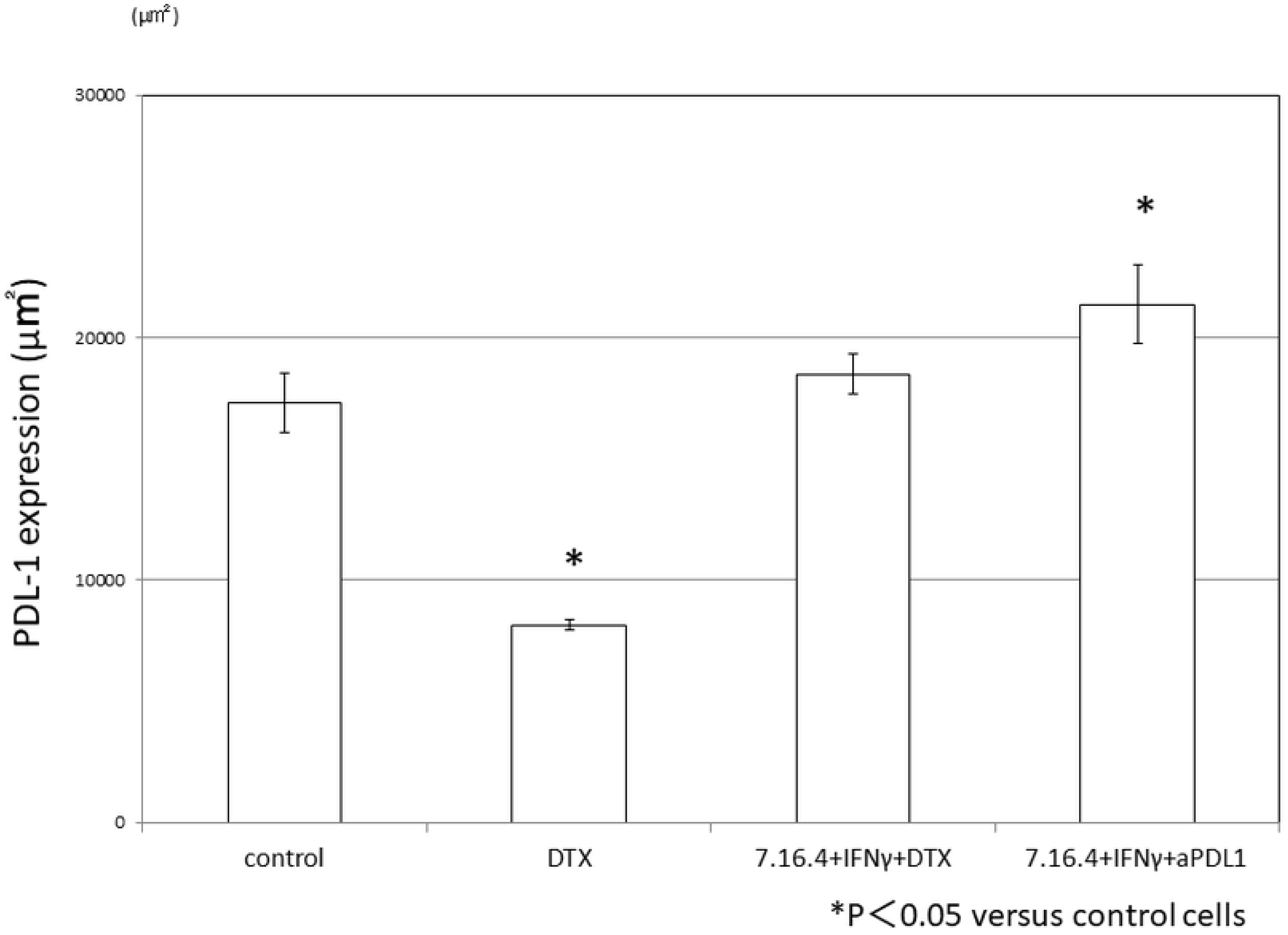

As shown in the images of immunofluorescence staining, compared with the control group, CD8 and expression was significantly higher in 7.16.4 + IFN-γ + aPD-L1 group (p<0.05) than the other groups. The Gr-1 expression was significantly lower in 7.16.4 + IFN-γ + aPD-L1 and 7.16.4 + IFN-γ + DTX group (p<0.05). In DTX alone group the PD-L1 expression significantly lower. And the PD-L1 expression in 7.16.4 + IFN-γ + PD-L1 group was significantly higher than the control group.

## Discussion

### The combined antibody + IFN-γ therapy directly acted on cancer cells and inhibited the cell proliferation against resistant cancer cell line

At first, we made the resistant cell line for the antibody therapy based on H2N113 cells derived from Balb / C mouse to maintain the same immunogenicity. We developed the resistant cancer cell line treatment by a method widely performed before ^[18]^.

The mechanism of drug resistance is not only the problem of signal transduction in cells, but also the problem related to the cells’ ability to avoid the immune system. It is a fairly complicated process that other reports have tried to elucidate ^[19] [20] [21]^.

We don’t know what kind of resistant mechanism is working for this resistant cell which we made this time, but we propose that there are various and complicated resistant mechanisms, similar to the clinical setting. ^[22] [18]^ We evaluated whether the combined anti-HER2 antibody and IFN-γ therapy shows antitumor effect in vitro, utilizing the resistant cell line (Figure1).

At first, anti-HER2 antibody therapy alone didn’t show any antitumor effect. That proved that the cancer cells had acquired resistance to antibody therapy. Combination therapy with IFN-γ significantly inhibited cell proliferation. In vitro experiment showed that the antibody and IFN-γ combined therapy has a direct effect on cancer cells by affecting cellular proliferation. The cell number was not decreased. This is consistent with our previous in vitro experiment before. ^[17]^

We previously demonstrated that the combined antibody and IFN-γ therapy changes the malignancy of cancer cells. ^[17]^

We also showed that this combination therapy acts on P27kip1 in RAJI and the antiCD20 antibody, inhibiting cellular proliferation. ^[23]^

Based on these prior results, we believe that the combined therapy would have an inhibitory effect on the malignant features of cancer cells, which is related to the problem of therapeutic resistance and cellular proliferation. There was no report about the interaction of the intracellular signal transduction by EGFR and EGFR2 with IFNGR and STAT1, but Yun et al reported the new opinion about interesting transduction that relate to antitumor effect of the antiHER2 antibody therapy. They showed that the activation of STAT1 by IFN-γ, which is secreted by immune cells, played an important role to diminish the intracellular signal transduction of Her2. ^[24]^

In this in vitro study, it was thought that the diminishment of the signal transduction was strongly induced by the combined Ab and IFN-γ therapy against the resistant cell line.

### The combination therapy for the resistant strain shows antitumor effect in the mouse Xenograft model

Next, we evaluated whether the combined anti-HER2 antibody and IFN-γ therapy showed antitumor effect even if it is in vivo experiment that used the same immunogenicity. The combined therapy significantly inhibited the growth of the tumor volume (Figure2). We thought that the antitumor immunity also worked in vivo, not only the direct action to the cancer cell which we showed in vitro.

### Combination therapy and CD8T cells (Cytotoxic T cell)

There have been many reports regarding the association between invasive ability of cytotoxic T cell to tumor tissue and treatment outcomes in many kinds of cancer ^[25] [26]^.

Many researchers have reported on the role of the tumor infiltrating lymphocytes. There are many reports that show positive correlations between CD8T cells and the prognosis, while there is reverse correlation between Th2 and Treg cells and prognosis.

The accumulation of CD8T cells in tumor tissue was significantly increased in the combined therapy group as compared to other groups using resistant cancer cells (Figure3).

Also, we showed that the positive correlation between antitumor effect and the number of infiltrating CD8T cells depends on the difference in drug treatment.

### About CD8T cells accumulation and MDSCs

It is well known that the tumor cells use various methods to escape from the immune system.

Treg cell or MDSCs are well known for immunosuppressive cells located in tumor tissue ^[20]^.

In the immunostaining of the tumor tissue, accumulation of MDSCs was significantly decreased in the combined-therapy group. This is consistent with our prior report, and it was shown that the secretion of cytokine from cancer tissue was inhibited by the combination therapy. Even though we didn’t prove the direct association between the increased accumulation of CD8T cells and decreased accumulation of MDSCs, we thought that the decrease of the accumulation of MDSCs contributed to the increase in the accumulation of CD8T cells.

Tumor-specific T cells, as part of adaptive immunity, presents a molecular MHC Class?on the cell surface and shows strong antitumor effect, through the entry of cancer antigenic peptide and co-stimulation. HER2 antibody and the stimulation by IFN-γ raises the MHC class I expression of APC’s, which leads us to believe that adaptive immunity is promoted by combination therapy.

### Combination therapy and ADCC activity and Natural killer cell

It is known that ADCC (antibody dependent cell cytotoxicity) plays an important role in the antitumor effect by the molecular target medicine ^[27] [28]^.

NK-cells, macrophages, and some neutrophils possess ADCC activity ^[28]^. Some researchers reported that IFN-γ often increases the Fc_γ_R expression of NK-cell. Motohashi and Taniguchi et al reported that IFN-γ from NK-cells reinforce the direct cell damaging action through NKT cells. IFN-γ generally works to enhance host immunity ^[29] [30]^.

However, when IFN-γ is used by monotherapy, tumor growth is actually increased in in vivo experiments. Therefore, IFN-γ also has a negative effect on host immunity. For example, it raises the PD-L1 expression of tumors ^[31]^. In fact, in our study, IFN-γ treatment raised the expression of PD-L1 in the tumor tissue (Figure7). Therefore, tumor size was larger than any other treatment. On the other hand, PD-L1 expression at the local site decreased in the combination therapy.

Also, Yanjun reported that there is an association between the CD8T cells with the tumor tissue of the patient with breast cancer and expression of PD-L1 and outcome ^[32]^.

### Combination therapy and NKT cells

Recently, some researchers have reported the contribution of NKT cells to antitumor immunity. For example, Motohashi et al. reported about the immunotherapy using NKT cells ^[29] [30] [33]^.

NKT cells also plays a role in the innate immunity, which doesn’t rely on the MHC system, allowing targeting of all cancer cells. The NKT cells strongly activate the above NK-cell and CD8T cells by secreting IFN-γ^[29] [30] [33]^.

We did not examine NKT cell’s accumulation in the tumor tissue this time. But we thought that the IFN-γ from NKT cells played an important role and showed the antitumor effect in this experiment. In the future, we hope to examine the contribution of NKT cells in the combination therapy in the future.

### Combination therapy and tumor suppressive MDSCs

Immunosuppressive cells, such as regulatory T cell (Treg) and myeloid derived suppressor cells(MDSCs), are very important for cancer progression. Because these cells inhibit CTL, NK-cell and NKT cells to accumulate into the tumor tissue. ^[34] [35]^

When we examined the difference between the tumor size and the accumulation of MDSCs this time, we showed a reverse correlation in the number of MDSCs and the volume of tumor. We showed that combination treatment using the molecular target drug and IFN-γ decreases the accumulation of MDSCs in cancer tissues, with an accumulation of CD8T cells as a result.

Even if cancer cells develop resistance to molecular targeting drugs, the combination therapy produces some chemokines involved in the accumulation of MDSCs. In the future, we should examine the changes of chemokines in the tumor microenviroment.

### About clinical application of the combination therapy with IFN-γ

Next, we evaluated which therapy would show the best antitumor effect in vivo, in order to determine if this combined therapy could be applied in the clinical setting. (Figure3).

The current recommended first-line pharmacotherapy for Her2-positive cancer is chemotherapy with antiHER2 antibody recommended ^[6] [9] [8] [36]^.

A similar regimen of chemotherapy is administered for breast cancer treatment.

When cancer cells acquire the pharmacotherapy-resistance, typically within one year, response to therapy halts ^[18]^.

The median overall survival in the GBG 26/BIG 3-05 phase III study of breast cancer is 24.9 months ^[37]^,

The progression free survival in the Toga study of gastric cancer is 6.7 months, and the median duration of overall survival is 13.8 months. With such poor prognosis, further improvement is required in the clinical setting [6].

When the progression was found during first line chemotherapy, we have an option to change the chemo drug or choose TDM-1, and various RCT is ongoing about the next choice now ^[38]^.

From our prior results, the 7.16.4+DOC therapy showed the antitumor effect that was equivalent to 7.16.4+ IFN-γ, and triple therapy with 7.16.4+DOC+ IFN-γ showed the highest antitumor effect. ^[17]^

We evaluated the next therapy for cancers that had progressed while on first-line chemotherapy. And we tested the immuno-check point inhibitor (CPI), which has attracted attention recently. We used the antiPD-L1 antibody for the immuno-check point inhibitor, because that showed more antitumor effect than antiPD-1 antibody in previous experiment.

The combination therapy utilizing 7.16.4+ IFN-γ and DOC or antiPD-L1 antibody treatment showed high antitumor effect. CD8T cells did not accumulate in 7.16.4+DOC+ IFN-γ therapy, while a significantly greater number of CD8T cells accumulated in the 7.16.4 + IFN-γ+ antiPD-L1 antibody.

We thought that the DTX has a direct cytotoxic effect, and antitumor immune effect by the host was worked in the without DTX therapy. We think that the immunotherapy is more useful for the patient with recurrent cancer than chemotherapy. Typically, the patients’physical condition is poor, due to recent chemotherapy use, in addition to the progression of the disease itself. We showed that antiHER2 antibody + IFN-γ + CPI showed a significantly higher anti-tumor effect than single agent treatment utilizing CPI alone.

## Conclusion

In this study, we proved that the combined antiHER2 antibody and IFN-γ therapy showed the antitumor effect for the cancer cell which had acquired resistance. We also showed that the tumor effect was higher than the current conventional therapy and that this combination therapy would be the best choice for many patients who suffer from cancer progression due to the development of treatment resistance.

